# Multicenter preclinical studies as an innovative method to enhance translation: a systematic review of published studies

**DOI:** 10.1101/591289

**Authors:** Victoria T. Hunniford, Agnes Grudniewicz, Dean A. Fergusson, Emma Grigor, Casey Lansdell, Manoj M. Lalu, on behalf of the Canadian Critical Care Translational Biology Group

## Abstract

Multicenter preclinical studies have been suggested as a method to improve reproducibility, generalizability and potential clinical translation of preclinical work. In these studies, multiple independent laboratories collaboratively conduct a research experiment using a shared protocol. The use of a multicenter design in preclinical experimentation is a recent approach and only a handful of preclinical multicenter studies have been published. Here, we systematically identify, assess and synthesize published preclinical multicenter studies investigating interventions using *in vivo* models. Synthesized data included study methods/design, basic characteristics, outcomes, and barriers and facilitators. Study risk of bias, completeness of reporting and the degree of collaboration were evaluated using established methods. The database searches identified 3095 citations and 12 studies met inclusion criteria. The multicenter study design was applied across a diverse range of diseases including stroke, heart attack, traumatic brain injury, and diabetes. The median number of centers was 4 (range 2-6) and the median sample size was 135 (range 23-384). Most studies had lower risk of bias and higher completeness of reporting than typically seen in single-centered studies. Only four of the twelve studies produced results consistent with previous single-center studies, highlighting a central concern of preclinical research: irreproducibility and poor generalizability of findings from single laboratories. Our review suggests that multicenter preclinical studies may provide a method to robustly assess therapies prior to considering clinical translation. Registered with PROSPERO CRD42018093986.

## Introduction

The translation process of preclinical findings into clinical practice is fraught with time lags, steep costs, and considerable failure rates [1–7]. It has been suggested that one of the problems contributing to translational failures lies outside of clinical research itself, and instead originates in the preclinical stage of research [8–10]. Translational barriers associated with preclinical research include poor study design and reporting that make reproducibility difficult; biased selection of animal models and small sample sizes which reduces inferential strength; and publication bias which may distort evidence and justification to proceed to first-in-human trials [5, 6, 11]. In order to increase the chance of ‘bench-to-bedside’ translation success, various measures to improve the state of preclinical research have been suggested [12, 13]. One measure is the application of multicenter experimentation in preclinical studies, analogous to what is commonly done in clinical trials [14, 15]. In both clinical and preclinical research, multicenter studies can assess external validity and inherently test reproducibility, while also increasing efficiency in meeting enrolment numbers [9]. In addition, rigorously designed and reported multicenter studies offer the opportunity to enhance internal validity and increase transparency [16].

To improve the process of translation, multiple calls from the biomedical science community have been made to adopt the multicenter preclinical approach [6, 12–14, 16, 17]. Some recent examples have been published that exemplify the successful implementation of this approach [18–20]. As interest in multicenter preclinical studies grows, and to demonstrate their value (if any), it is imperative that a systematic evaluation should be performed of the studies conducted to date. This will inform and optimize future multicenter preclinical studies by identifying knowledge gaps and producing an evidence map of current practices and outcomes [5–7].

The objective of this systematic review was to identify and qualitatively summarize the preclinical multicenter study literature. All multicenter *in vivo* preclinical studies of interventions were included. We compared and contrasted the methods and organization of these experiments. Quality of reporting, risk of bias, degree of collaboration, and barriers/enablers to multicenter study conduct were assessed. Finally, we considered how results of these studies and the use of the multicenter study design informed the translation of biomedical research.

## Results

### Search results and study characteristics

The database searches identified a total of 3095 papers after duplicates were removed (Fig 1). Two additional papers were identified through a search of references of included papers. After title, abstract, and full-text screening twelve articles met eligibility criteria (Tables 1 and 2).

**Fig 1.**
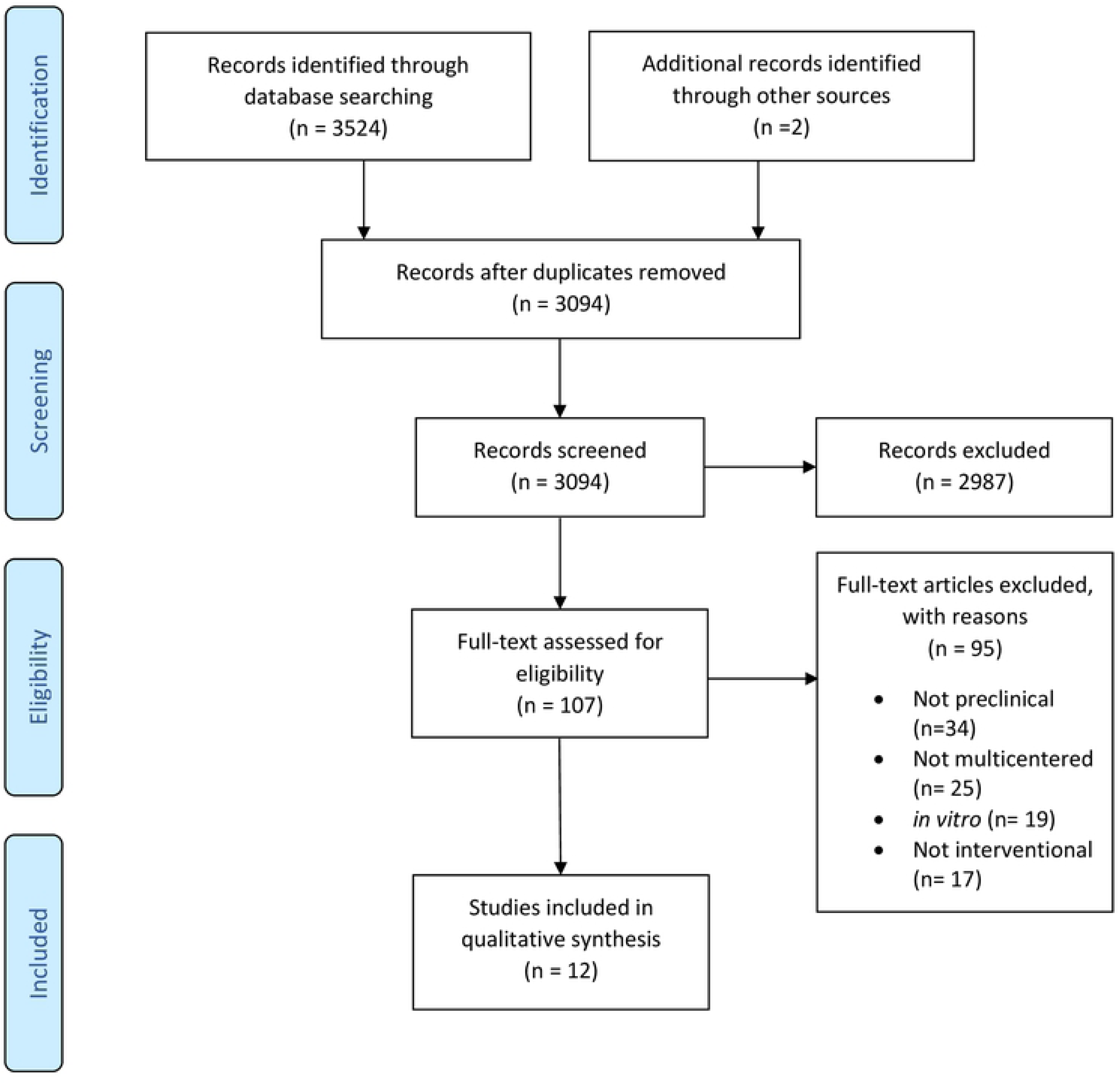
Preferred reporting items for systematic reviews and meta-analysis (PRISMA) flow diagram for study selection.

**Table 1.**
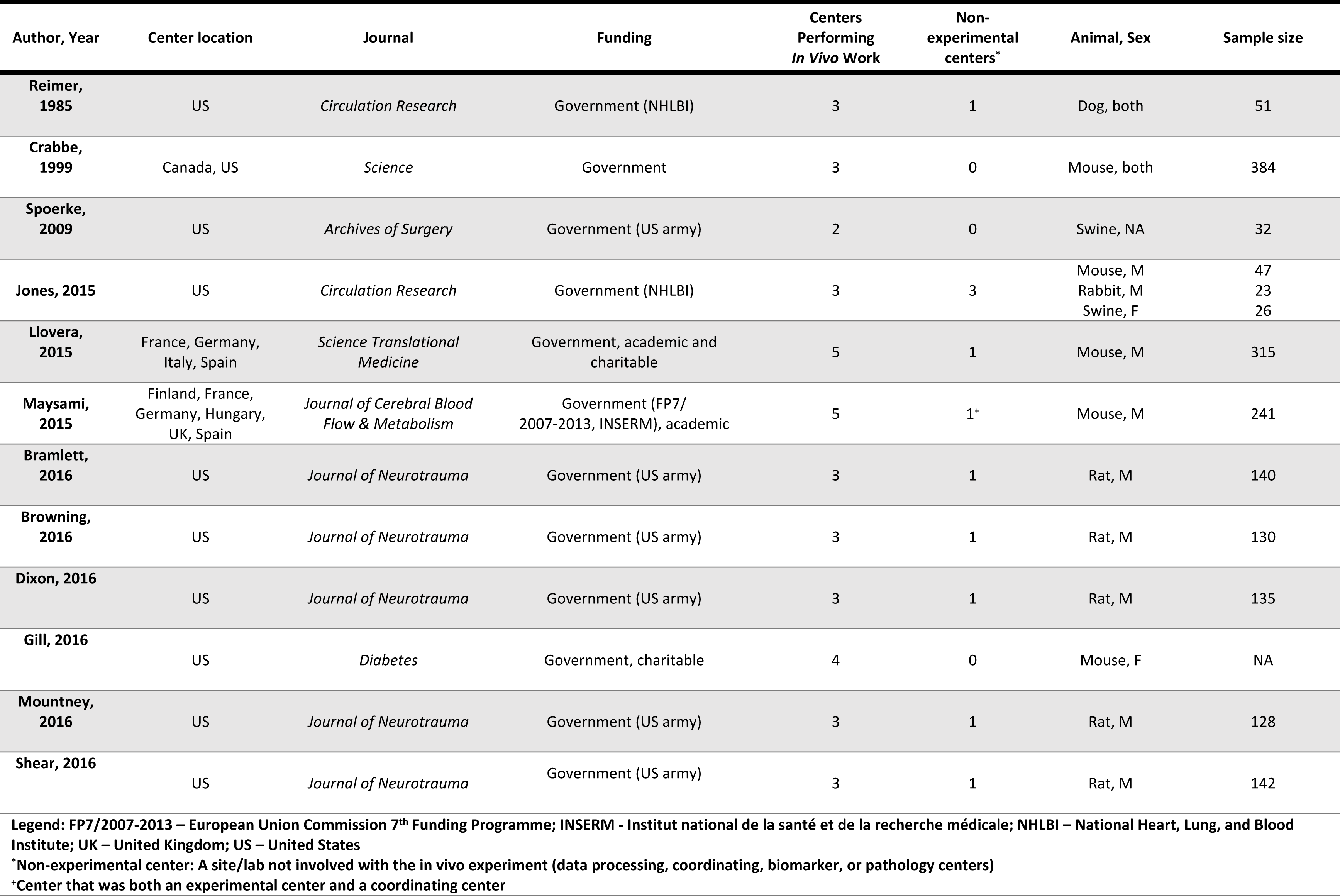
Basic study characteristics of preclinical multicenter studies.

| Author, Year | Center location | Journal | Funding | Centers Performing In Vivo Work | Non-experimental centers* | Animal, Sex | Sample size |
| --- | --- | --- | --- | --- | --- | --- | --- |
| Reimer, 1985 | US | *Circulation Research* | Government (NHLBI) | 3 | 1 | Dog, both | 51 |
| Crabbe, 1999 | Canada, US | *Science* | Government | 3 | 0 | Mouse, both | 384 |
| Spoerke, 2009 | US | *Archives of Surgery* | Government (US army) | 2 | 0 | Swine, NA | 32 |
| Jones, 2015 | US | *Circulation Research* | Government (NHLBI) | 3 | 3 | Mouse, M<br>Rabbit, M<br>Swine, F | 47<br>23<br>26 |
| Llovera, 2015 | France, Germany, Italy, Spain | *Science Translational Medicine* | Government, academic and charitable | 5 | 1 | Mouse, M | 315 |
| Maysami, 2015 | Finland, France, Germany, Hungary, UK, Spain | *Journal of Cerebral Blood Flow & Metabolism* | Government (FP7/2007-2013, INSERM), academic | 5 | 1<sup>+</sup> | Mouse, M | 241 |
| Bramlett, 2016 | US | *Journal of Neurotrauma* | Government (US army) | 3 | 1 | Rat, M | 140 |
| Browning, 2016 | US | *Journal of Neurotrauma* | Government (US army) | 3 | 1 | Rat, M | 130 |
| Dixon, 2016 | US | *Journal of Neurotrauma* | Government (US army) | 3 | 1 | Rat, M | 135 |
| Gill, 2016 | US | *Diabetes* | Government, charitable | 4 | 0 | Mouse, F | NA |
| Mountney, 2016 | US | *Journal of Neurotrauma* | Government (US army) | 3 | 1 | Rat, M | 128 |
| Shear, 2016 | US | *Journal of Neurotrauma* | Government (US army) | 3 | 1 | Rat, M | 142 |

Legend: FP7/2007-2013 – European Union Commission 7<sup>th</sup> Funding Programme; INSERM - Institut national de la santé et de la recherche médicale; NHLBI – National Heart, Lung, and Blood Institute; UK – United Kingdom; US – United States
\*Non-experimental center: A site/lab not involved with the in vivo experiment (data processing, coordinating, biomarker, or pathology centers)
\*Center that was both an experimental center and a coordinating center

**Table 2.**
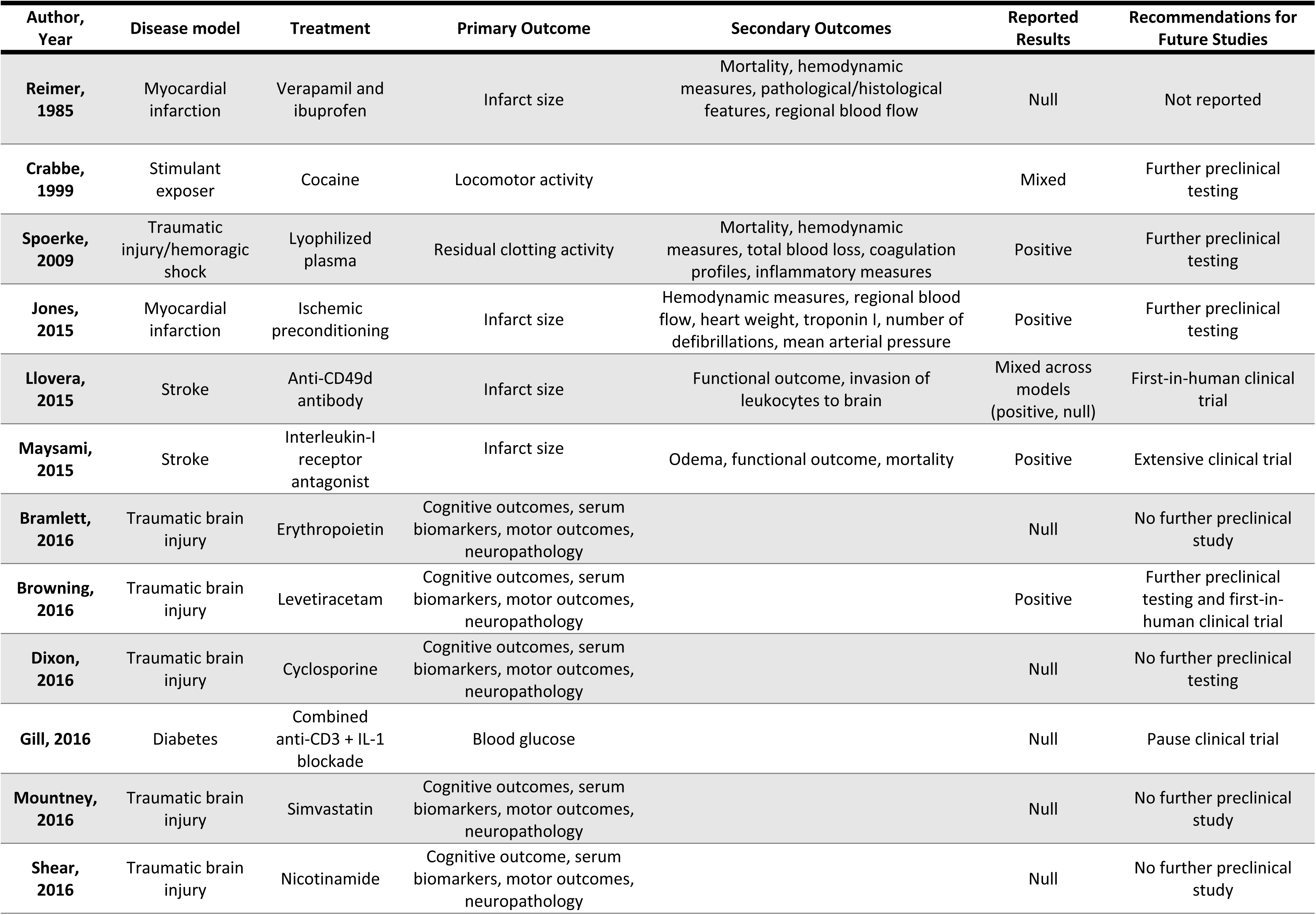
Study design characteristics of preclinical multicenter studies.

The identified studies fell into six clinical domains: traumatic brain injury (*n* = 5), myocardial infarction (*n* = 2), stroke *(n* = 2), diabetes (*n* = 1), traumatic injury (*n* = 1), and effects of stimulate exposure (*n* = 1). Nine of twelve studies were published in 2015 and 2016, three studies were published each in 1985, 1999, and 2009. Ten studies were fully funded by government sources; two studies had partial funding from government and charitable or academic funding.

Three studies were international (studies with centers located in the USA, Germany, France, Canada, Finland, Hungary, Italy, the United Kingdom, and Spain), and nine studies were conducted solely in the USA (all centers located in the USA). The median number of total centers involved per multicenter study was 4 (range: 2-6), and the median number of experimental centers performing *in vivo* work was 3 (range: 2-5). Nine studies (75%) reported having non-experimental centers involved, such as a coordinating center, data processing center, biomarker core, and a pathology core. Five different species of animals were used by the studies: mice (*n* = 5), rats (*n* = 5), swine (*n* = 2), rabbits (*n* = 1), and dogs (*n* = 1). The median sample size was 135 (range 23-384 animals), and a total 1794 animals were used across the twelve studies, 93% of which were lab rodents (mice and rats).

### Reported outcomes

Four of the studies (33%) reported that the treatment showed statistically significant, positive results; six studies reported that the treatment showed non-significant or null results; two studies reported that the results were mixed (positive and null) across different animal models of the disease of interest [18] or outcome measures [21] (Table 2). Based on their respective results, eleven studies made explicit statements of recommendations, or future directions for the therapy of study. Five studies stated that they would conduct further testing or recommended that further preclinical testing be done on the tested therapy. Four studies indicated they would not continue testing or recommended that no further preclinical testing be done. Two studies recommended to proceed with first-in-human clinical trials, and one recommended conducting an extensive clinical trial. Three of the studies that recommended further preclinical testing had mixed (*n* = 1) or positive (*n* = 2) results; the three studies that recommended to proceed with human clinical trials had mixed (*n* = 1) and positive results (*n* = 2), and the five studies that suggested that there should be no further testing (clinical or preclinical) all had null results (Fig 2). Brief synopses of the twelve studies can be found in supporting information (S1 Text), along with sample statements of their future recommendations (S2 Table).

**Fig 2.**
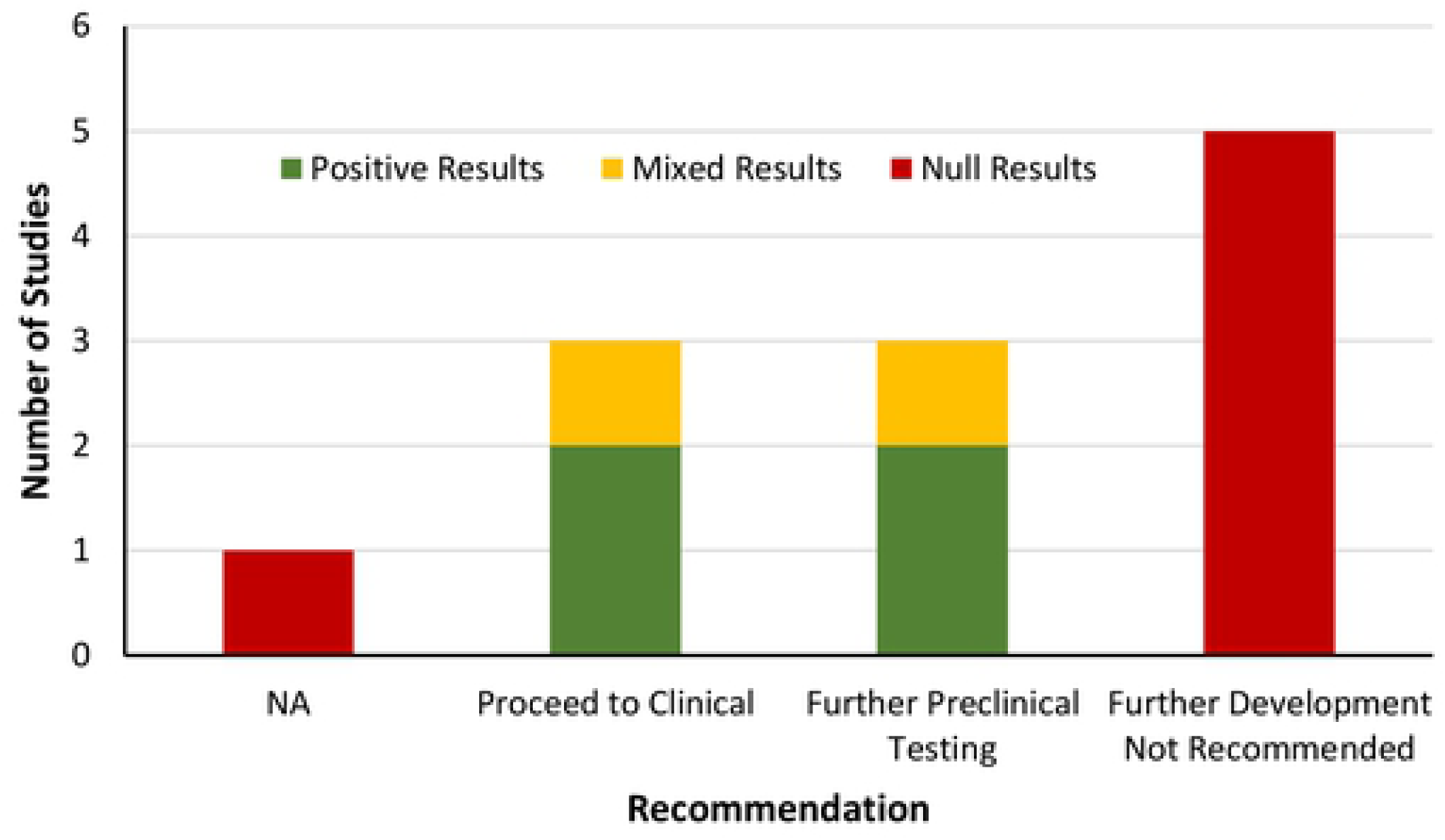
Preclinical multicenter studies reported recommendations for future studies along with the reported results of each study. The recommendation to proceed to clinical includes recommending first-in-human trials and extensive clinical trials.

### Risk of Bias

None of the 12 studies (0%) were considered low risk of bias across all ten domains (Table 3). Ten studies randomized animals to experimental groups and two of these reported the method of random sequence generation. Nine studies had a low risk of detection bias by blinding of outcome assessors. Eight studies were at low risk of performance bias by blinding personnel administering interventions. All but one study was unclear if animals were randomly housed during the experiments. Five studies from the same research consortium (Operation Brain Trauma Therapy) had high risk of bias for other sources of bias due to potential industry-related influences (Table 3). The four ‘other sources’ of risk of bias assessments for each study is found in the supporting information (S3 Table).

**Table 3.** Risk of bias assessment of multicenter preclinical interventional studies.

| Study | Sequence generation* | Baseline characteristics | Allocation concealment* | Random housing | Blinding of personnel | Random outcome assessment | Blinding of outcome assessment | Incomplete outcome data* | Selective outcome reporting | Other sources of bias** |
| --- | --- | --- | --- | --- | --- | --- | --- | --- | --- | --- |
| Reimer, 1985 | U<sup>1</sup> | U | L | U | H<sup>3</sup> | U | L | U | L | U |
| Crabbe, 1999 | U<sup>1</sup> | L | U | L | U | U | U | L | L | H |
| Spoerke, 2009 | U<sup>1</sup> | L | U | U | U | L | U | U | L | U |
| Jones, 2015 | L | L | U | U | L | L | L | L | L | L |
| Llovera, 2015 | L | L | U | U | L | L | L | L | L | L |
| Maysami, 2015 | H<sup>2</sup> | H | L | U | L | L | L | U | H | H |
| Bramlett, 2016 | U<sup>1</sup> | U | U | U | L | L | L | U | L | H |
| Browning, 2016 | U<sup>1</sup> | U | U | U | L | L | L | U | L | H |
| Dixon, 2016 | U<sup>1</sup> | U | U | U | L | L | L | U | L | H |
| Gill, 2016 | H | L | U | U | U | L | U | U | U | L |
| Mountney, 2016 | U<sup>1</sup> | U | U | U | L | L | L | U | L | H |
| Shear, 2016 | U<sup>1</sup> | U | U | U | L | L | L | U | L | H |

**Legend:** H = High risk of bias, L = Low risk of bias, U = Unclear risk of bias
\*Items in agreement with the Cochrane Risk of Bias tool [22]
**Baseline Characteristics:** Low risk = Relevant baseline characteristics equal between experimental groups or controlled for. Unclear = Relevant baseline characteristics unreported. High risk = Relevant baseline characteristics unbalanced between experimental groups and not controlled.
**Random Housing:** Low risk = Animal cages were randomly placed within an animal room/facility, Unclear = Housing placement unreported, High risk = Animals place in non-random arrangement in animal room/facility.
**Blinding of Outcome Assessment:** Low risk = Outcome assessors were blinded to the study groups when assessing endpoints/animals Unclear = Insufficient information to determine if outcome assessors were blinded during assessment. High Risk = Outcome assessors not blinded to the study groups.
**Incomplete Outcome Data:** Low risk = N values were consistent between methods and results for the outcomes. Unclear = N value was either not presented in the methods or in the results, and therefore there is insufficient information to permit judgement. High risk = N values were not consistent between methods and results for the outcomes.
**Selective Reporting:** Low risk = The methods section indicated pre-specified outcome measure. Unclear: Was not clear about the pre-specified primary endpoints and outcome results. High risk = The outcome was presented in the results but not pre-specified in the methods section.
\* other sources include funding influences, conflicts of interest, contamination, unit of analysis errors
<sup>1</sup>Method of randomization not specified

### Reporting quality

Overall, completeness of reporting across all twelve studies was high, ranging from 62% to 100% of checklist items being reported. One of the twelve studies reported on all the 29 items in the preclinical multicenter reporting checklist. The domains with the highest completeness of reporting included replicates (biological vs. technical), statistics, blinding, and discussion (Table 4). The domains for standards, randomization, sample size estimation, and inclusion/exclusion criteria were variable in the completeness of reporting. The introduction and abstract domain had the lowest completeness of reporting, as eleven of the twelve studies did not report that the study was multicentered in the title (item 1, Table 4) - though four of the eleven studies used a synonym for multicenter (i.e. consortium or cross-laboratory), and less than half indicated the number of participating centers in the abstract. Though it was not an item in the reporting checklist, four studies included *preclinical* in the title, two studies had *animal (or swine) model* in the title, and five studies from the same traumatic brain injury consortium did not indicate that the study was preclinical (or synonym) in the title. Additional papers [23, 24] that accompanied the five traumatic brain injury studies included *preclinical* in the paper title. Reporting assessment for all twenty-nine items across the twelve studies can be found in the supporting information (S4 Table).

**Table 4.** Frequency of reported preclinical multicenter checklist items.

| Domain | # | Item Description | % of studies that reported |
| --- | --- | --- | --- |
| Intro/<br>abstract | 1 | Identification as a multicenter study in title | 8 |
|  | 2 | Abstract states number of participating centers | 42 |
| Standards | 3 | Community based reporting guidelines listed | 17 |
|  | 4 | Names of each participating center listed | 100 |
|  | 5 | List roles of participating centers (central coordinating center, experimental site) | 83 |
|  | 6 | No changes, or if applicable major changes to study protocol after commencement are documented | 100 |
| Replicates<br>(biological vs.<br>technical) | 7 | Results substantiated by repetition under a range of conditions at each site | 100 |
|  | 8 | Number of subjects per outcome | 100 |
|  | 9 | Number of measurements per subject for one experimental outcome stated | 83 |
|  | 10 | Number of subjects per center | 75 |
| Statistics | 11 | List of the total number of subjects used in each experimental group | 83 |
|  | 12 | List of all statistical tests used | 100 |
|  | 13 | Definition of the measure of central tendency | 100 |
|  | 14 | Definition of the measure of dispersion | 100 |
| Randomization | 15 | Random group assignment reported | 100 |
|  | 16 | Description of the method of random group assignment | 33 |
| Blinding | 17 | Experimenters blinded to group allocation during conduct of the experiment | 75 |
|  | 18 | Experimenters blinded to group allocation during result assessment | 83 |
| Sample Size<br>Estimation | 19 | Description of an *a priori* primary outcome | 100 |
|  | 20 | Sample size for each site computed during study design | 33 |
|  | 21 | Description of the method of sample size determination | 33 |
| Inclusion and<br>Exclusion<br>Criteria | 22 | Total number of animals for the experiment reported | 50 |
|  | 23 | Description of the criteria used for the exclusion of any data or subjects | 42 |
|  | 24 | List losses and exclusions of animals at the end of experiment | 50 |
|  | 25 | All outcomes described, or description of any outcomes that were measured and not reported in the results section | 100 |
|  | 26 | Previous or pilot/preliminary studies performed and listed | 83 |
|  | 27 | Results were significant, or if not, null or negative outcomes included in the results | 100 |
| Discussion | 28 | Limitations of the study are documented | 83 |
|  | 29 | Discrepancies in results across centers expected or absent, or if not, they discussed | 100 |

**Legend:** Coloured cells indicate the frequency (%) of item reported over all included studies. Frequency (%) ranges: 0-33 = red; 34-74 = yellow; 75-100 = green.

### Degree of collaboration

Overall, the twelve studies scored medium to high in the degree of collaboration (Table 5). The ‘development’ domain of collaboration had the greatest number of studies given ‘high’ degree of collaboration assessment. The ‘execution’ and ‘coordination’ domains had similar overall degree of collaboration assessments, though there was one more study assessed as ‘low’ in the coordination domain, and one more study assessed as ‘high’ in the execution domain. One study [19] had high degree of collaboration across all three domains, and one study received an unclear score of 0 across all three domains [25] – meaning that this study was unclear in the reporting for each domain of collaboration.

**Table 5.** Degree of collaboration assessment.

| Author, Year | Development | Execution | Coordination |
| --- | --- | --- | --- |
| Reimer, 1985 | 2 | 2 | 3 |
| Crabbe, 1999 | 3 | 3 | 2 |
| Spoerke, 2009 | 0 | 0 | 0 |
| Jones, 2015 | 3 | 3 | 3 |
| Llovera, 2015 | 2 | 3 | 3 |
| Maysami, 2015 | 2 | 1 | 1 |
| Bramlett, 2016 | 3 | 2 | 2 |
| Browning, 2016 | 3 | 2 | 2 |
| Dixon, 2016 | 3 | 2 | 2 |
| Gill, 2016 | 2 | 3 | 1 |
| Mountney, 2016 | 3 | 2 | 2 |
| Shear, 2016 | 3 | 2 | 2 |

### Reported barriers and facilitators

Five of the twelve studies (42%) explicitly reported on the barriers and facilitators to conducting a multicenter study. The most frequently reported barrier identified in all five studies [18–20, 26, 27] was the establishment of a consistent protocol, with attention to exact experimental details across research centers. In addition to the challenge of the initial protocol development, studies reported difficulty in centers strictly adhering to the established protocol throughout the entirety of the study. One study [20] had considerable issues in adhering to the protocol, and in effect had to modify their methods through the course of the study.

Three studies [18–20] reported differences in equipment and resources across centers as a barrier that made it difficult to conduct a collaborative project and to communicate what measurements and endpoints would be assessed. Specific experimental conditions that investigators were unable or unwilling to modify included animal models of the disease, animal housing conditions, the separate labs’ operating and measurement procedures, equipment, and institutional regulations. There was also inconsistent funding across research centers. Different centers had separate budgets with different amounts of funding that could be allocated to the study. If the protocol was to be harmonized, then it had to be adapted to fit each center’s budget accordingly (i.e., the center with the smallest budget set the spending limit) [20]. Alternatively, centers developed a general protocol but adapted it to fit their own respective budget with what resources they had. Another barrier identified was ethics approval for animal experiments at all the centers [18]. This was especially significant when centers were located in multiple countries, as each country had different regulations for ethical approval [18, 20].

Jones *et al.* [19] suggested that a clearly defined experimental method was facilitated by employing a pilot test through all the centers and subsequently developing a protocol collaboratively. Developing a defined experimental protocol also included establishing an agreed upon timeline, laboratory setup and method of analysis and measurement. Maysami *et al.* and Reimer *et al.* [20, 26], retrospectively suggested that a similar approach might have enhanced the conduct of both of their studies. Another study reported that the facilitator was the use of a centralized center for administration and data processing [18, 19]. The validity of reports depends on statistical and data management control; and having one center coordinate these operations reduces the chances of error or bias in the analysis. Other facilitators were related to the interpersonal aspect of collaboration. These included having investigator leadership through regular conferences and check-ins from beginning to end of the project [28] and building upon previously established personal/professional relationships between investigators [19].

## Discussion

Multiple calls for the use of and reviews on the topic of multicenter study design in preclinical research have been published [5–7, 17, 23]. Here, we have moved beyond narrative pieces to synthesizing characteristics and outcomes from all preclinical multicenter studies conducted to date. Our results suggest that this is an emerging, novel and promising area of research, with twelve studies conducted to date and the majority being published since 2015. Notable differences between the studies included clinical spheres, number of centers involved, sample sizes, and degrees of collaboration. Similarities between studies included sources of funding (largely governmental), countries involved (approximately 80% of centers were located in the USA), and the species of animal used (93% lab rodents).

A noteworthy finding is the discordance of results between previous single-center studies and subsequent multicenter preclinical studies. Four studies reported that their results confirmed previous single-center findings, six found no effect, and two found mixed effects. An explanation for why half of the multicenter studies reported null results, when the smaller primary studies conducted at single centers reported positive results, may reflect the increased sample size, methodological rigor, routine oversight and quality control of these multicenter studies. All of these factors would inherently lead to a more precise evaluation of the intervention’s effects. A similar trend has been noted in clinical studies with decreased interventional effects, as studies move from single to multicenter studies [29]. Our findings allude to one of the central contentious issues of preclinical research: irreproducibility and poor generalizability of findings from single laboratories. Based on the studies’ results and conclusions, eleven of the twelve studies made explicit recommendations or suggestions for future studies and development. This observation enhances the rationale behind the use of a multicentric design in preclinical research. As interventions are considered for translation, shifting away from traditional small-scale single-centered studies to a more methodologically rigorous, multicentric design may ultimately reduce translational failures and research waste.

Previous preclinical systematic reviews of single center studies have found unclear or high risk of bias in most domains [30, 31]. In contrast, we found that multicenter studies generally had a lower risk of bias than typical single-centered studies. It has been established that practices such as blinding and randomization are important aspects of study methodology known to affect the internal validity of both preclinical and clinical studies. Most studies had low risk of bias in the three domains directly related to issues of randomization and blinding. Failing to use these methods represents a lack of rigor and is one of the suggested reasons behind failed translation [8].

Overall, the included studies had high completeness of reporting across most domains. Many of the items that all or most studies reported on were a part of the domains that address statistics and replicates. The items that studies reported at a lower frequency were specific to multicenter designs, such as indicating the number of participating centers in the abstract and identifying as a multicenter study in the title. One potential explanation for this finding is that guidelines and standards for multicenter studies are just emerging, and there has yet to be any reporting recommendations specific to a preclinical multicenter design. Nonetheless, compared to previous reporting assessments of single-centered preclinical studies [32, 33], the completeness of reporting for the twelve preclinical multicenter studies was appreciably higher.

In general, we found a medium to high degree of collaboration. The domain of protocol development had the greatest proportion of studies assessed as high degree of collaboration, with no studies assessed as ‘low.’ Based on the degree of collaboration criteria, this means that in most studies the protocol was developed *a priori* by all centers with or without a pilot study. Protocol execution had markedly lower scores, suggesting that centers may have found it difficult to maintain a high degree of collaboration when the protocol was being executed.

### Recommendations

After reviewing the barriers and facilitators reported in included studies, as well as preforming an assessment of the completeness of reporting, we provide recommendations for future preclinical multicenter studies in Box 1.

#### Box 1 Recommendations for preclinical multicenter studies

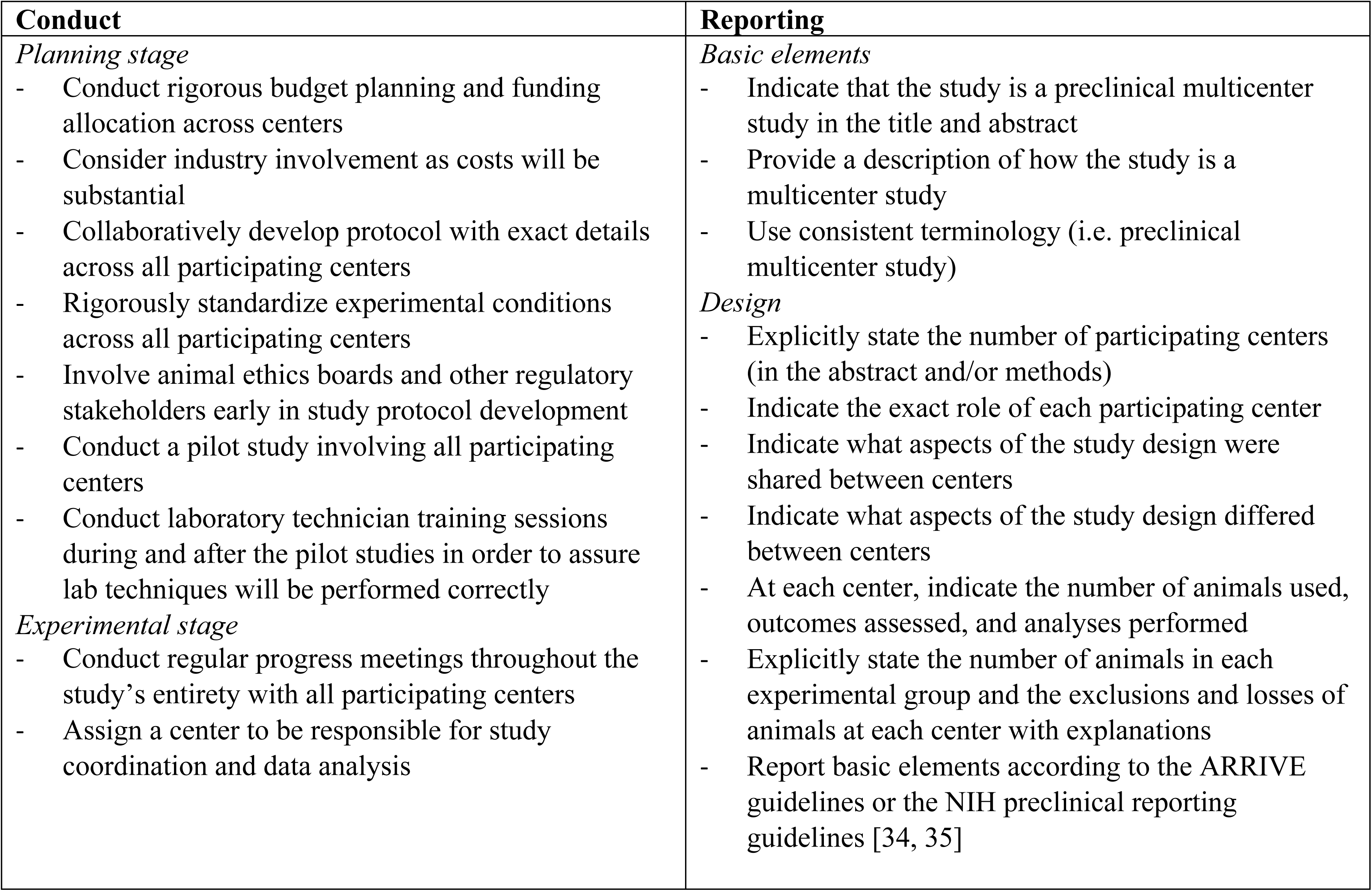

| Conduct | Reporting |
| --- | --- |
| *Planning stage*<br>- Conduct rigorous budget planning and funding allocation across centers<br>- Consider industry involvement as costs will be substantial<br>- Collaboratively develop protocol with exact details across all participating centers<br>- Rigorously standardize experimental conditions across all participating centers<br>- Involve animal ethics boards and other regulatory stakeholders early in study protocol development<br>- Conduct a pilot study involving all participating centers<br>- Conduct laboratory technician training sessions during and after the pilot studies in order to assure lab techniques will be performed correctly<br>*Experimental stage*<br>- Conduct regular progress meetings throughout the study’s entirety with all participating centers<br>- Assign a center to be responsible for study coordination and data analysis | *Basic elements*<br>- Indicate that the study is a preclinical multicenter study in the title and abstract<br>- Provide a description of how the study is a multicenter study<br>- Use consistent terminology (i.e. preclinical multicenter study)<br>*Design*<br>- Explicitly state the number of participating centers (in the abstract and/or methods)<br>- Indicate the exact role of each participating center<br>- Indicate what aspects of the study design were shared between centers<br>- Indicate what aspects of the study design differed between centers<br>- At each center, indicate the number of animals used, outcomes assessed, and analyses performed<br>- Explicitly state the number of animals in each experimental group and the exclusions and losses of animals at each center with explanations<br>- Report basic elements according to the ARRIVE guidelines or the NIH preclinical reporting guidelines [34, 35] |

### Strengths and limitations

A strength of this review is the systematic synthesis of the published literature, which allows for a more in-depth assessment of the state of this field of research. However, the scope of this review could also be seen as a limitation, as no quantitative results from the individual studies were assessed due to the heterogeneity of the studies. Nonetheless, we were able to broadly describe and compare results between these multicenter studies and preceding single-center investigations. The application of rigorous inclusion criteria limited the eligible studies to interventional, controlled-comparison studies, which could omit valuable qualitative information that may have come from the excluded studies of non-controlled and/or observational designs. However, it should be noted that we purposely chose these criteria in order to understand how these studies may inform translation to human patients.

### Conclusion

Despite there only being 12 multicenter studies published to date, this review demonstrates the value of a multicentric study design in preclinical research, as it was found that these studies employed extensive quality control and study oversight. This potentially led to the observed lower risk of bias and higher completeness of reporting versus typical single laboratory studies. Moreover, we believe the identified barriers and facilitators, and the list of core recommendations for preclinical multicenter studies, may optimize future applications of this approach. Importantly, this review provides evidence that preclinical multicenter studies may be used as a tool to inform further development and future studies of promising clinical therapies at the confirmatory, preclinical stage of translational research.

## Material and Methods

This systematic review was reported in accordance with the Preferred Reporting Items for Systematic Review and Meta-Analysis (PRISMA) guidelines [36]. A copy of the PRISMA checklist is provided in the supporting information (S5 Checklist). The protocol was registered with the International Prospective Register of Systematic Reviews (PROSPERO CRD42018093986) and was posted on the Collaborative Approach to Meta-Analysis and Review of Animal Data from Experimental Studies (CAMARADES) website (http://syrf.org.uk/protocols/).

### Eligibility Criteria

#### Population

The population of interest was preclinical, interventional, multicenter, controlled-comparison studies. *Preclinical* was defined as research that is medically relevant and is conducted using nonhuman models prior to being tested in human subjects. *Multicenter* was defined as cooperative research formally conducted between multiple research centers (sites). Models were limited to *in vivo* experiments but were not limited by the clinical scope or domain of the preclinical study. Furthermore, selected studies were not limited by the individual studies’ type of intervention, protocol, variables of interest or results.

#### Intervention, comparators, outcomes

There were no limitations to specific intervention, comparator or outcomes of individual studies included.

#### Study design

Eligible preclinical studies included *in vivo*, controlled comparison, interventional studies of randomized and non-randomized designs. *In vivo* experiments needed to be conducted at two or more independent sites for the study to qualify as multicentric. The sites needed to also share more than just general study objectives to be considered multicentered. Features that met the multicenter criteria included, but were not limited to: shared design, protocol, animal model, method of analysis, primary endpoints tested with or without identical measurement apparatuses; separate centers for coordination, protocol development, and data analysis; and study objective, timelines, protocols, and dissemination strategies developed *a priori*. Veterinary clinical trials, *in vitro* and *ex vivo* studies (with no *in vivo* component), and retrospective data analysis from multiple sites were excluded.

#### Search strategy

The search strategy was created in collaboration with an information specialist (Risa Shorr, The Ottawa Hospital) and the investigators. Electronic databases Embase (Embase Classic and Embase 1947 – 30 January 2018), and MEDLINE (Ovid MEDLINE Epub Ahead of Print, In-Process & Other Non-Indexed Citations, Ovid Medline Daily and Ovid Medline 1946 to 29 January 2018) were searched on January 31, 2018. The search strategy was developed using keywords related to multicenter studies as well as preclinical research, such as preclinical OR animal model experimental model, AND multicenter OR cross laboratory. An updated search was run on April 24th, 2018 using a modified search strategy in the same databases as the first search. No study scope, date or language limits were imposed. The search strategy was tested through the inclusion of six target articles [17–21, 26] identified by the investigators prior to the systematic search. A second, independent librarian peer reviewed the search strategy according to the Peer Review of Electronic Search Strategy (PRESS) framework [37]. The updated search strategy is presented in the supporting information (S6 Appendix), as well as the PRESS review (S7 Appendix).

#### Screening and data extraction

The results from the literature search were uploaded to Distiller Systematic Review Software (DistillerSR®; Evidence Partners, Ottawa, Canada). DistillerSR is a cloud-based program that facilitates the review process by managing studies through customized screening, auditing and reporting. Duplicate references were removed and two reviewers (VTH and CL/EM) independently screened titles and abstracts based on the eligibility criteria. Any disagreements were resolved by consensus. For the second stage of screening, two reviewers (VTH and MML) independently screened the full text reports of included references based on the eligibility criteria. Disagreements were solved via consensus.

Data was extracted using a standardized extraction form developed in DistillerSR that was piloted by the primary reviewer (VTH) on five studies. The extraction form was revised based on feedback from a senior team member (MML) and was then made into an Excel spreadsheet where extracted data was saved. Data included characteristics of the studies: publication details (authors, year published, journal), the country(ies) where the study was conducted, sources of funding, the number of centers involved (experimental and non-experimental), the disease model, animal species and sex, sample size, treatment/exposure, primary and secondary study outcomes, the reported results, statements of barriers and facilitators, and statements of recommendations and suggestions for future testing of the specific therapy being investigated.

#### Assessing completeness of reporting and risk of bias

Risk of bias and quality of reporting were assessed independently by two reviewers (VTH and MML), and disagreements were resolved via consensus. All randomized, interventional studies were assessed as high, low, or unclear for the 10 domains of bias adapted from the SYRCLE “Risk of Bias” assessment tool for preclinical *in vivo* studies [38]. The ‘other sources’ of risk of bias domain was divided into 4 sub-domains (funding influences, conflicts of interest, contamination, and unit of analysis errors). An overall ‘other’ risk of bias assessment was given based on the following: overall high risk of bias if one or more of the four other sources were assessed as high; overall unclear risk of bias if two or more of the four other sources were assessed as unclear; and overall low risk of bias if three of the four other sources were assessed as low.

Quality of reporting of the studies was assessed using a checklist modified from various sources: consolidated Standards of Reporting Trials (CONSORT) [39]; the National Institutes of Health (NIH)’s principles and guideline for reporting preclinical research [35]; and the Good Clinical Practice (GCP) Guidance Document: E6(R2) [40]. The checklist is provided in the supporting information (S8 Table) with details on the sources for each item.

#### Assessing degree of collaboration

An assessment of the degree of collaboration reported was made for each multicenter study. The degree of collaboration assessment criteria developed by the investigators was based on a validated scale used to measure collaboration among grant partners. The original scale developed by Frey and colleagues used five levels of collaboration ranging from 1 to 5, and a score or 0 for no interaction at all [41]. Assessments were given based on the reporting of (or lack thereof) study design elements in 3 domains: protocol development, protocol execution, and collaboration. These elements have been suggested to contribute to success in multicenter clinical, and early preclinical experiments, such as the use of separate centers for data processing and coordination, a collaborative development and shared protocol between all centers involved, standardized reporting across all centers involved, etc. [5, 6, 42, 43]. The degree of collaboration in each domain was assessed with a four-point scale (0-3) defined as low, medium, high, or unclear and scored from 1 (for low) to 3 (for high). Domains that were unclear were given a score of 0. The summary table used color to visualize the degree of collaboration across the three domains (red = low, yellow = medium, green = high, white = unclear) (S9 Table). Degree of collaboration was assessed independently by two reviewers (VTH and MML).

#### Results and data synthesis

The results from the systematic review were extracted descriptively and data was interpreted through a narrative synthesis. The findings and methods of the systematic review were reported in accordance with the PRISMA guidelines [36, 44]. Descriptive data was synthesized and presented through tabulation of textual elements [45]. Studies were arranged in tables based on study design and basic characteristics, degree of collaboration assessments, and risk of bias assessments. A synthesis of any statements and examples pertaining to barriers and facilitators in conducting a multicenter study was also performed on the included multicenter research studies. Given that the focus of this review is reporting design and characteristics rather than a specific outcome (and the heterogenous nature of preclinical study topics) performing a meta-analysis was not appropriate.

#### Deviations from protocol

The original protocol submitted to PROSPERO indicated that the degree of collaboration would be evaluated through a summed score of all three domains, resulting in a numerical value ranging from 0 to 9. After expert feedback, it was decided not to use a quantitative score and to assess the domains individually. We elected to assign colors for each assessment to provide a visual representation of the degree of collaboration across the three domains.

## Contributions (using CRediT Taxonomy)

Guarantor: MML; Conceptualization: VTH, MML, DAF; Methodology: VTH, MML, DAF; Investigation: VTH, CL, EG; Writing – original draft: VTH; Writing – review and editing: all authors; Supervision: MML, AG, and DAF; Resources: MML.

## Acknowledgments

VTH was supported by a Queen Elizabeth II Graduate Scholarship in Science and Technology and MML is supported by The Ottawa Hospital Anesthesia Alternate Funds Association and the Scholarship Protected Time Program, Department of Anesthesiology and Pain Medicine, uOttawa. We would like to thank Risa Shorr (Information specialist, The Ottawa Hospital) for providing assistance with the generation of a systematic search strategy and article retrieval. We would also like to Dr. Alison Fox-Robichaud from the Canadian Critical Care Translational Biology Group for providing critical feedback on the manuscript.

## Supporting Information

### S1 Text Study Overviews

1. In a study by Reimer et al. (1985) [26], three independent laboratories collaborated to develop models to test potential ischemic myocardium protection therapies, using two standardized, well-characterized canine models of myocardial infarction. Using the two different dog models (conscious model of coronary occlusion, and unconscious model of 3-hour ischemia in open-chest), the researchers tested the effects of verapamil and ibuprofen (therapies) on infarct size. The pooled results from all three centers demonstrated that neither drug limited infarct size in either model. It was later published that the participating laboratories discovered through statistical and hard evidence that a forth participating lab initially involved in the study had fraudulent data, in the sense that data had been completely fabricated by the lead researcher at one of the centers [46]. The data from this lab was not included in the multicenter study paper. The detection of the fraudulent data would not have been possible if not for the design of a multicenter study. The fraud was detected by the large discrepancies of study outcome data between to offending center and the other centers involved in the study. It was later confirmed by the coordinating as they performed own investigation into the lead scientist of the offending center.
2. Crabbe et al. (1999) [21] performed a large study across 3 laboratories. The main objective was to test the behavioural variability in mice of different genetic strains, sexes, and laboratory environments. The evaluation was done with identical testing apparatus and protocol across all three labs. The potentially clinically relevant portion of this study was an assessment of cocaine’s effect on behavior (i.e. locomotor activity). The study found that cocaine effects on locomotor activity had a strong relationship with genetic differences on the laboratory giving the tests but was negligible for sex differences and source of mice (i.e. shipped from a supplier or were bred locally).
3. Spoerke et al. (2009) [25] tested whether lyophilized plasma (LP) is as safe and effective as fresh frozen plasma (FFP) for resuscitation after severe trauma. They used a swine model of severe injury across animal laboratories of two level I trauma centers, to test the lyophilized plasma for factor levels and clotting activity before lyophilization and after reconstitution. The swine model was developed and performed at one of the centers and was learned and performed at a second center to test for reproducibility. They found that LP decreased clotting factor activity and was equal to FFP in terms of efficacy.
4. Jones et al. (2015) [19] aimed to develop a multicenter, randomized controlled clinical-like infrastructure for preclinical evaluation of cardioprotective therapies using mice, rabbit and pig models. The researchers established the Consortium for preclinical assessment of cardioprotective therapies – called CAESAR, to test the ability of ischemic preconditioning (IPC) to reduce infarct size of a myocardial infarction. IPC involved short episodes of blood restriction to the heart – which is an experimental technique for producing resistance to blood loss. It has demonstrated to activate the largest number of protective pathways and is currently the most reproducible cardioprotection intervention to date. Six centers (2 centers/animal model) tested the therapy in the three animal models with shared protocols, and found the results were similar across centers and that IPC significantly reduced infarct size in all three species.
5. Llovera and colleagues (2015) [18] performed a preclinical randomized controlled multicenter trial (pRCT) to test the potential of Anti-CD49d antibodies as treatment from acute brain ischemia. These antibodies have shown promise as a form of therapy in individual laboratories by inhibiting the migration of leukocytes into the brain following acute brain lesion. Leukocyte invasion is known as brain inflammation and is a key mechanism that mediates secondary neuronal injury after a stroke. Six independent European research centers centrally coordinated a study to test the antibody using mouse models of stroke, and the pooled results demonstrated that the antibody significantly reduced leukocyte invasion after mechanically induced stroke (distal occlusion of the middle cerebral artery). They found that the treatment did not limit infarct size and concluded that this therapy should not be evaluated further in clinical trials.
6. Maysami and colleagues (2015) [20] conducted a cross-laboratory study in five centers (4 research, 1 coordinating) to test and interleukin receptor antagonist as a drug therapy for stroke. The antagonist drug is a naturally occurring drug that has been reported as beneficial for stroke recovery, as it inhibits the neuronal binding of the cytokine interleukin – a key mediator in neuronal injury (inflammation). The coordinating center developed and distributed the standard operating procedure to all centers, where mice were used as the model and ischaemia was induced both by permanent and transient occlusion. Drug effects on stroke outcome was evaluated by various means: lesion volume, oedema, neurological deficit scoring and post-treatment mortality. The results across all centers strongly support the therapeutic potential of the cytokine receptor antagonist in experimental stroke.
7. Five separate studies [47–51] that were coordinated by Operation Brain Trauma Therapy (OBTT) consortium [23, 24, 28]. Three independent centers collaborated to screen 5 different drugs that had been proposed for acute therapies in severe traumatic brain injury (TBI). The consortium was supported by the United States Army and had an overall approach of testing promising therapies in three-well established models of TBI in rats with a rigorous design. The end goal of the consortium was to test the 5 initial therapies in rats prior to considering further testing in a swine model of TBI. Based on the results, four of the five drugs preformed below or well below what was expected based on the previously published literature. It was reported that only levetiracetam would advance to testing in the swine model, and that an additional three drugs were being tested by the consortium as well.
8. Gill et al. (2016) [27] assessed the efficacy of combined anti-CD3 plus interleukin-1 blockade to reverse new-onset autoimmune diabetes in non-obese diabetic (NOD) mice. Their consortium was established by the National Institutes of Health Immune Tolerance Network and the Juvenile Diabetes Research Foundation. Four academic centers shared models and operating procedures, in nine different treatment groups. They found that the combined antibody treatment did not show reversal of diabetes across all sites. They did however conclude that intercenter reproducibility is possible with the NOD mouse model of diabetes.

**S2 Table.**
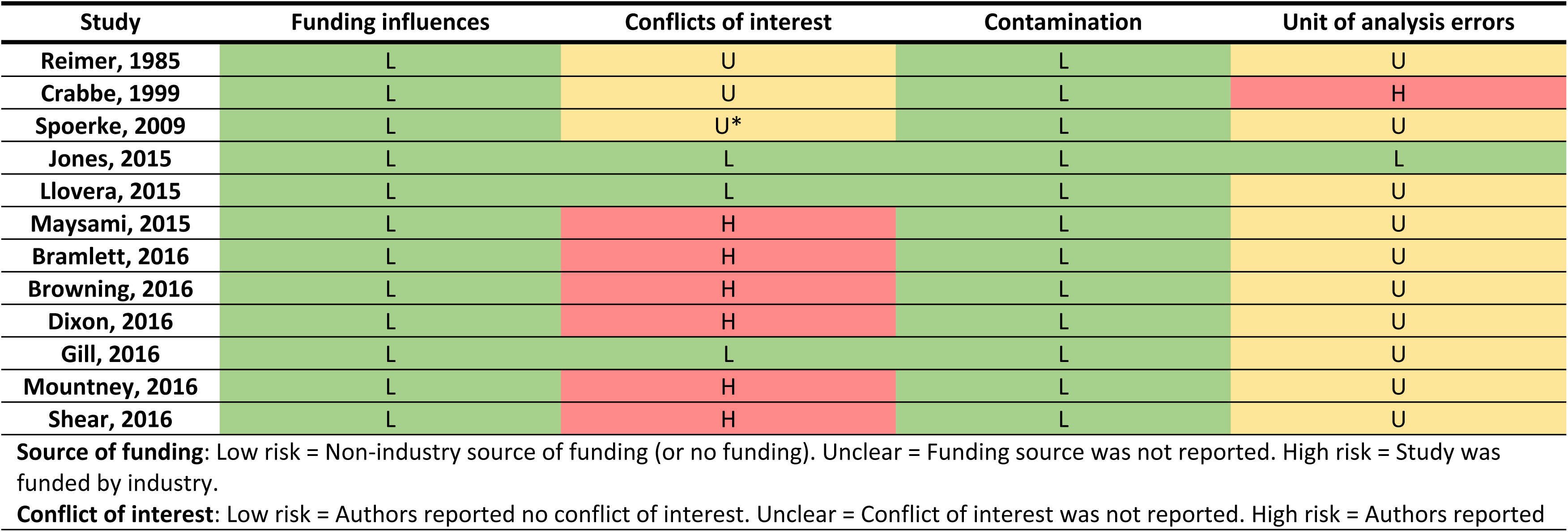
Risk of bias assessment for other sources of bias

**S3 Table.**
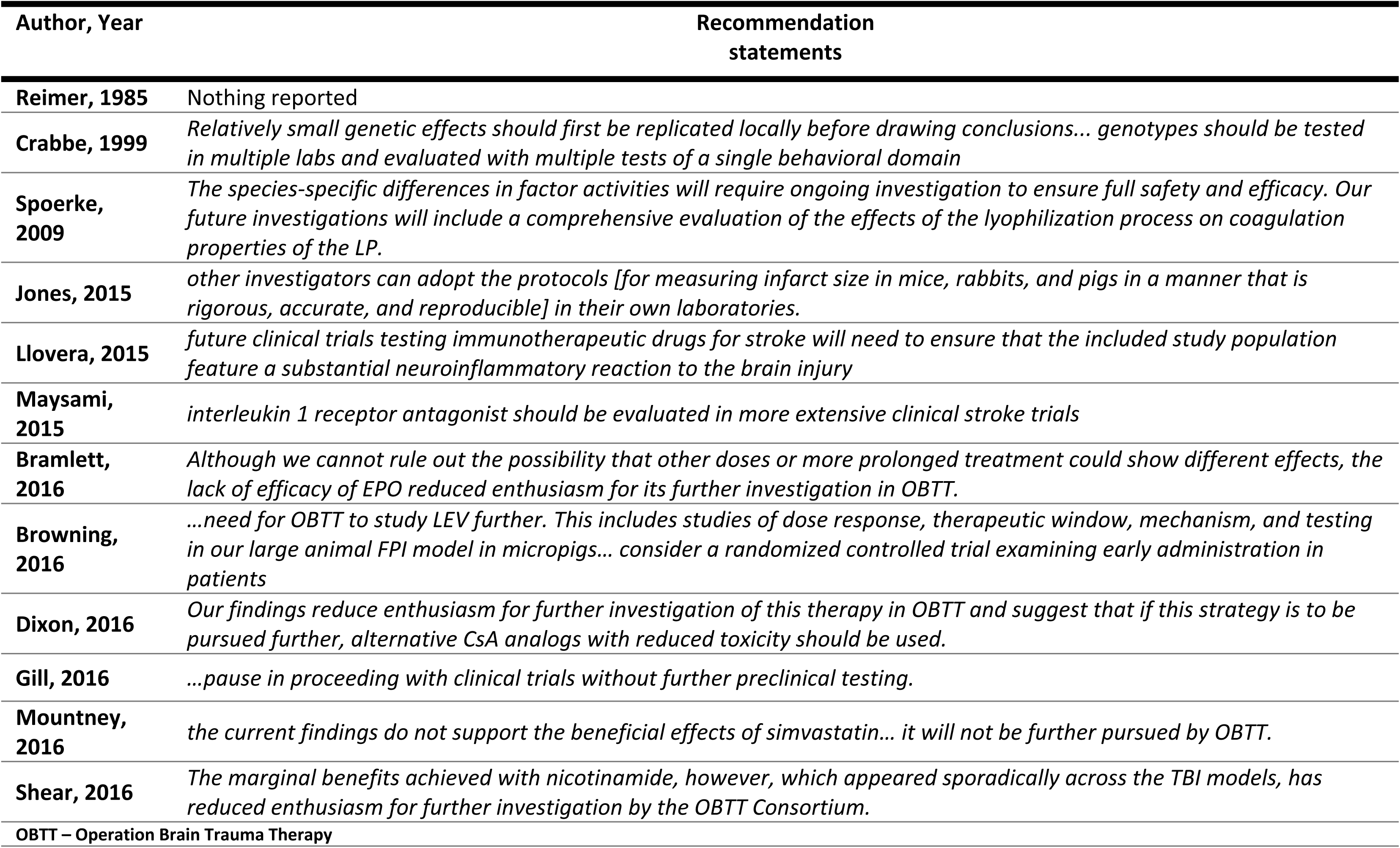
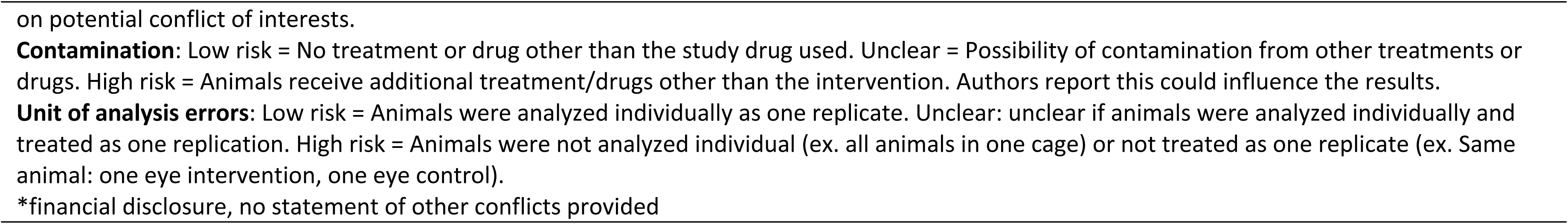
Statements of future recommendations

**S4 Table.**
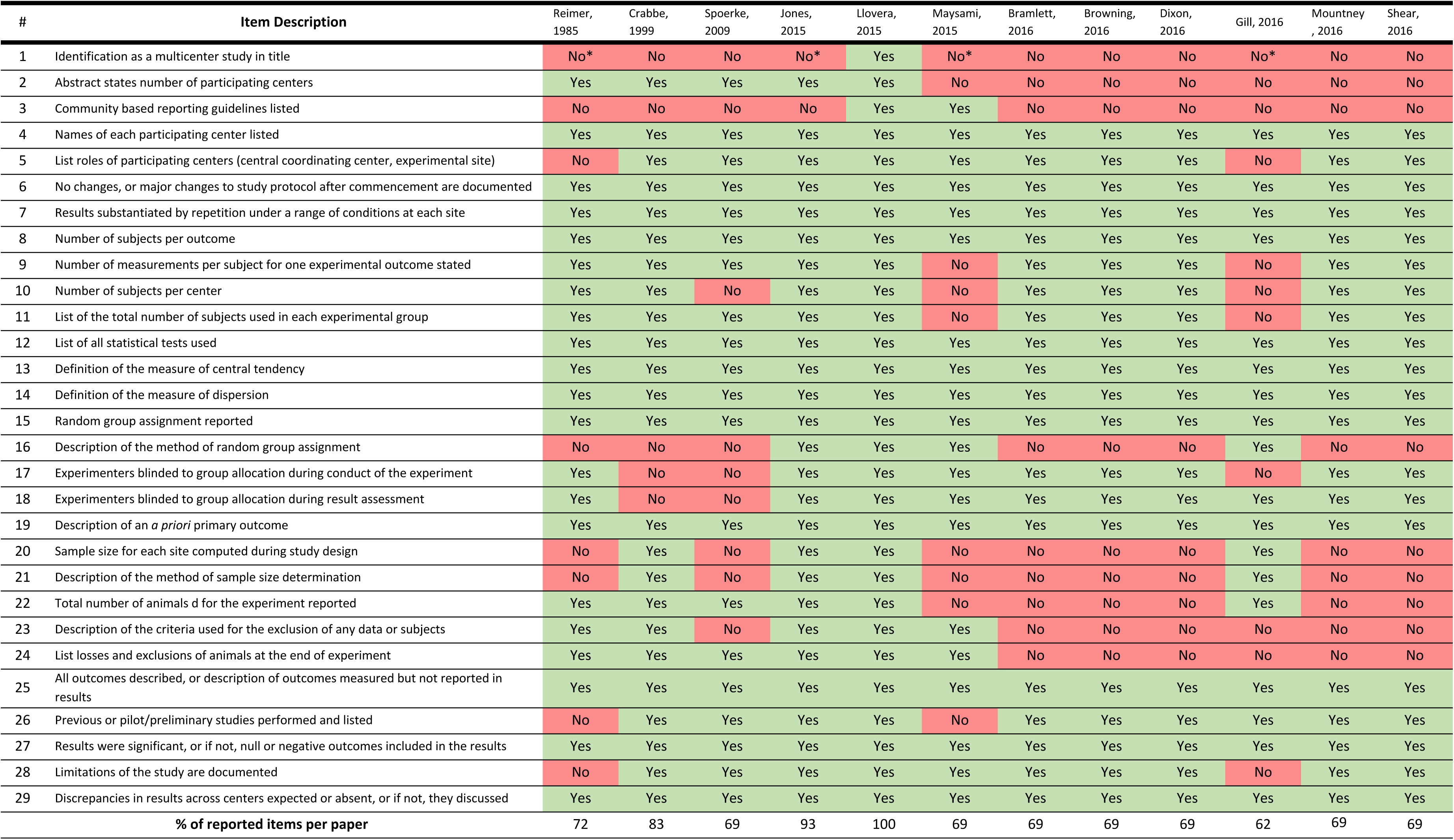
Completeness of reporting evaluation for 12 studies across 29 items

### S5 Checklist PRISMA Checklist

From: Moher D, Liberati A, Tetzlaff J, Altman DG, The PRISMA Group (2009). Preferred Reporting Items for Systematic Reviews and Meta-Analyses: The PRISMA Statement. PLoS Med 6(7): e1000097. doi:10.1371/journal.pmed1000097

| Section/topic | # | Checklist item | Reported on page # |
| --- | --- | --- | --- |
| **TITLE** |  |  |  |
| Title | 1 | Identify the report as a systematic review, meta-analysis, or both. | 1 |
| **ABSTRACT** |  |  |  |
| Structured summary | 2 | Provide a structured summary including, as applicable: background; objectives; data sources; study eligibility criteria, participants, and interventions; study appraisal and synthesis methods; results; limitations; conclusions and implications of key findings; systematic review registration number. | 2 |
| **INTRODUCTION** |  |  |  |
| Rationale | 3 | Describe the rationale for the review in the context of what is already known. | 3 |
| Objectives | 4 | Provide an explicit statement of questions being addressed with reference to participants, interventions, comparisons, outcomes, and study design (PICOS). | 4 |
| **METHODS** |  |  |  |
| Protocol and registration | 5 | Indicate if a review protocol exists, if and where it can be accessed (e.g., Web address), and, if available, provide registration information including registration number. | 1 & 21 |
| Eligibility criteria | 6 | Specify study characteristics (e.g., PICOS, length of follow-up) and report characteristics (e.g., years considered, language, publication status) used as criteria for eligibility, giving rationale. | 21-22 |
| Information sources | 7 | Describe all information sources (e.g., databases with dates of coverage, contact with study authors to identify additional studies) in the search and date last searched. | 22-23, 37 |
| Search | 8 | Present full electronic search strategy for at least one database, including any limits used, such that it could be repeated. | 22-23, 37 |
| Study selection | 9 | State the process for selecting studies (i.e., screening, eligibility, included in systematic review, and, if applicable, included in the meta-analysis). | 23 |
| Data collection process | 10 | Describe method of data extraction from reports (e.g., piloted forms, independently, in duplicate) and any processes for obtaining and confirming data from investigators. | 23-25 |
| Data items | 11 | List and define all variables for which data were sought (e.g., PICOS, funding sources) and any assumptions and simplifications made. | 23 |
| Risk of bias in individual studies | 12 | Describe methods used for assessing risk of bias of individual studies (including specification of whether this was done at the study or outcome level), and how this information is to be used in any data synthesis. | 24 |
| Summary measures | 13 | State the principal summary measures (e.g., risk ratio, difference in means). | 25-26 |
| Synthesis of results | 14 | Describe the methods of handling data and combining results of studies, if done, including measures of consistency (e.g., I<sup>2</sup>) for each meta-analysis. | 25-26 |
| Risk of bias across studies | 15 | Specify any assessment of risk of bias that may affect the cumulative evidence (e.g., publication bias, selective reporting within studies). | N/A |
| Additional analyses | 16 | Describe methods of additional analyses (e.g., sensitivity or subgroup analyses, meta-regression), if done, indicating which were pre-specified. | N/A |
| **RESULTS** |  |  |  |
| Study selection | 17 | Give numbers of studies screened, assessed for eligibility, and included in the review, with reasons for exclusions at each stage, ideally with a flow diagram. | 4-5 |
| Study characteristics | 18 | For each study, present characteristics for which data were extracted (e.g., study size, PICOS, follow-up period) and provide the citations. | 6-9 |
| Risk of bias within studies | 19 | Present data on risk of bias of each study and, if available, any outcome level assessment (see item 12). | 10-11 |
| Results of individual studies | 20 | For all outcomes considered (benefits or harms), present, for each study: (a) simple summary data for each intervention group (b) effect estimates and confidence intervals, ideally with a forest plot. | 4-16 |
| Synthesis of results | 21 | Present results of each meta-analysis done, including confidence intervals and measures of consistency. | N/A |
| Risk of bias across studies | 22 | Present results of any assessment of risk of bias across studies (see Item 15). | N/A |
| Additional analysis | 23 | Give results of additional analyses, if done (e.g., sensitivity or subgroup analyses, meta-regression [see Item 16]). | N/A |
| **DISCUSSION** |  |  |  |
| Summary of evidence | 24 | Summarize the main findings including the strength of evidence for each main outcome; consider their relevance to key groups (e.g., healthcare providers, users, and policy makers). | 16-20 |
| Limitations | 25 | Discuss limitations at study and outcome level (e.g., risk of bias), and at review-level (e.g., incomplete retrieval of identified research, reporting bias). | 19-20 |
| Conclusions | 26 | Provide a general interpretation of the results in the context of other evidence, and implications for future research. | 20 |
| **FUNDING** |  |  |  |
| Funding | 27 | Describe sources of funding for the systematic review and other support (e.g., supply of data); role of funders for the systematic review. | 29 |

## S6 Appendix Systematic search strategy

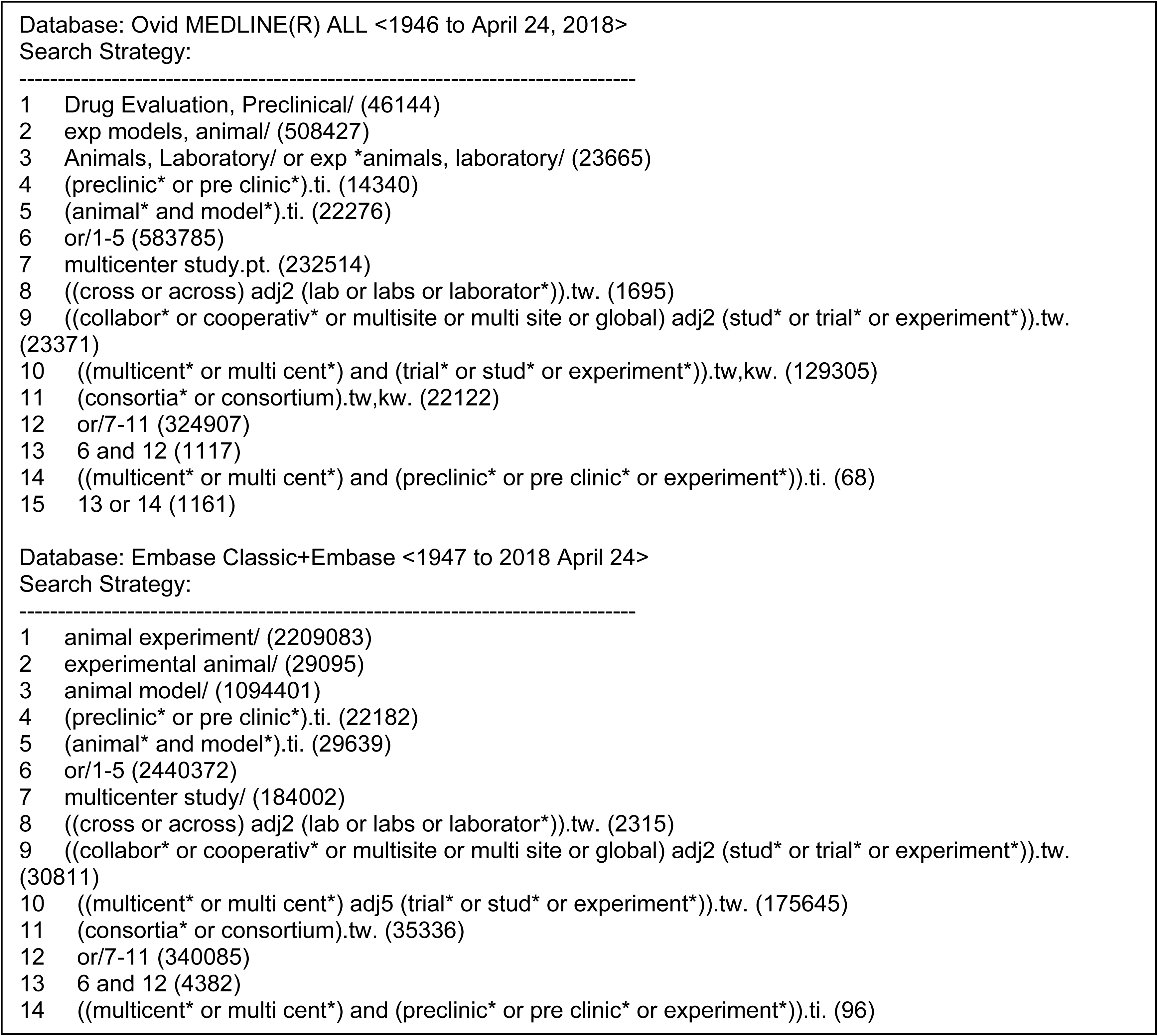

## S7 Appendix PRESS review of search strategy

***PRESS Guideline* — Search Submission & Peer Review Assessment**

**SEARCH SUBMISSION: THIS SECTION TO BE FILLED IN BY THE SEARCHER**

Searcher’s Name: Risa Shorr

Date Submitted: 2018-01-172018-01-17

Date Needed By: Click or tap to enter a date.

**Systematic Review Title**

Preclinical Multicenter Studies

**This search strategy is: (Highlight the appropriate response)**

A. My PRIMARY (core) database strategy — First time submitting a strategy for search question and database
B. My PRIMARY (core) strategy — Follow-up review NOT the first time submitting a strategy for search question and database. If this is a response to peer review, itemize the changes made to the review suggestions
C. SECONDARY search strategy— First time submitting a strategy for search question and database
D. SECONDARY search strategy – NOT the first time submitting a strategy for search question and database. If this is a response to peer review, itemize the changes made to the review suggestions.

**Database (ie, Medline, Cinahl**

Medline

**Interface (Ovid, Ebsco)**

Ovid

**Research Question. Describe the purpose of the search**

To summarise the literature on preclinical multicenter studies

**PICO Format (Outline the PICOs for your question – ie. Patient, Intervention, Comparison, Outcome and Study Design – as applicable)**

|  |  |
| --- | --- |
| P | Preclinical studies |
| I | Multicenter |
| C |  |
| O |  |

**Inclusion Criteria (List criteria such as age groups, study designs, etc, to be included)**

**[optional]**

**Exclusion Criteria (List criteria such as study designs, date limits, etc., to be excluded)**

***[optional]***

**Was a search filter applied?**

NoNo

**If YES, which one(s) (e.g., Cochrane RCT filter, PubMed Clinical Queries filter)? Provide the source if this is a published filter. [mandatory if YES to previous question — textbox]**

Other notes or comments you feel would be useful for the peer reviewer? [optional] Here are some target articles. All are captured with the search strategy.

Searching all animal groups and study type is too huge (~10000 refs in Medline) and then not all studies are indexed as multicenter.

**1. A cross-laboratory preclinical study on the effectiveness of interleukin-1 receptor antagonist in stroke.**

Maysami S; Wong R; Pradillo JM; Denes A; Dhungana H; Malm T; Koistinaho J; Orset C; Rahman M; Rubio M; Schwaninger M; Vivien D; Bath PM; Rothwell NJ; Allan SM.

Journal of Cerebral Blood Flow & Metabolism. 36(3):596-605, 2016 Mar. [Journal Article. Research Support, Non-U.S. Gov’t]

UI: 26661169

**2. Results of a preclinical randomized controlled multicenter trial (pRCT): Anti-CD49d treatment for acute brain ischemia.**

Llovera G; Hofmann K; Roth S; Salas-Perdomo A; Ferrer-Ferrer M; Perego C; Zanier ER; Mamrak U; Rex A; Party H; Agin V; Fauchon C; Orset C; Haelewyn B; De Simoni MG; Dirnagl U; Grittner U; Planas AM; Plesnila N; Vivien D; Liesz A.

Science Translational Medicine. 7(299):299ra121, 2015 Aug 05.

[Journal Article. Multicenter Study. Research Support, Non-U.S. Gov’t]

UI: 26246166

**3. The NHLBI-sponsored Consortium for preclinicAl assESsment of cARdioprotective therapies (CAESAR): a new paradigm for rigorous, accurate, and reproducible evaluation of putative infarct-sparing interventions in mice, rabbits, and pigs.**

Jones SP; Tang XL; Guo Y; Steenbergen C; Lefer DJ; Kukreja RC; Kong M; Li Q; Bhushan S; Zhu X; Du J; Nong Y; Stowers HL; Kondo K; Hunt GN; Goodchild TT; Orr A; Chang CC; Ockaili R; Salloum FN; Bolli R.

Circulation Research. 116(4):572-86, 2015 Feb 13.

[Journal Article. Multicenter Study. Research Support, N.I.H., Extramural]

UI: 25499773

**4. Different data from different labs: lessons from studies of gene-environment interaction. [Review] [83 refs]**

Wahlsten D; Metten P; Phillips TJ; Boehm SL 2nd; Burkhart-Kasch S; Dorow J; Doerksen S; Downing C; Fogarty J; Rodd-Henricks K; Hen R; McKinnon CS; Merrill CM; Nolte C; Schalomon M; Schlumbohm JP; Sibert JR; Wenger CD; Dudek BC; Crabbe JC.

Journal of Neurobiology. 54(1):283-311, 2003 Jan.

[Comparative Study. Journal Article. Research Support, Non-U.S. Gov’t. Research Support, U.S. Gov’t, Non-P.H.S.. Research Support, U.S. Gov’t, P.H.S.. Review]

UI: 12486710

**5. Genetics of mouse behavior: interactions with laboratory environment.**

Crabbe JC; Wahlsten D; Dudek BC.

Science. 284(5420):1670-2, 1999 Jun 04.

[Comparative Study. Journal Article. Research Support, Non-U.S. Gov’t. Research Support, U.S. Gov’t, Non-P.H.S.. Research Support, U.S. Gov’t, P.H.S.]

UI: 10356397

**6. Animal models for protecting ischemic myocardium: results of the NHLBI Cooperative Study. Comparison of unconscious and conscious dog models.**

Reimer KA; Jennings RB; Cobb FR; Murdock RH; Greenfield JC Jr; Becker LC; Bulkley BH; Hutchins GM; Schwartz RP Jr; Bailey KR; et al.

Circulation Research. 56(5):651-65, 1985 May.

[Comparative Study. Journal Article. Research Support, U.S. Gov’t, P.H.S.]

UI: 3838923

**Please copy and paste your search strategy here, exactly as run, including the number of hits per line. [mandatory]**

Database: Ovid MEDLINE(R) ALL <1946 to January 16, 2018>

Search Strategy:

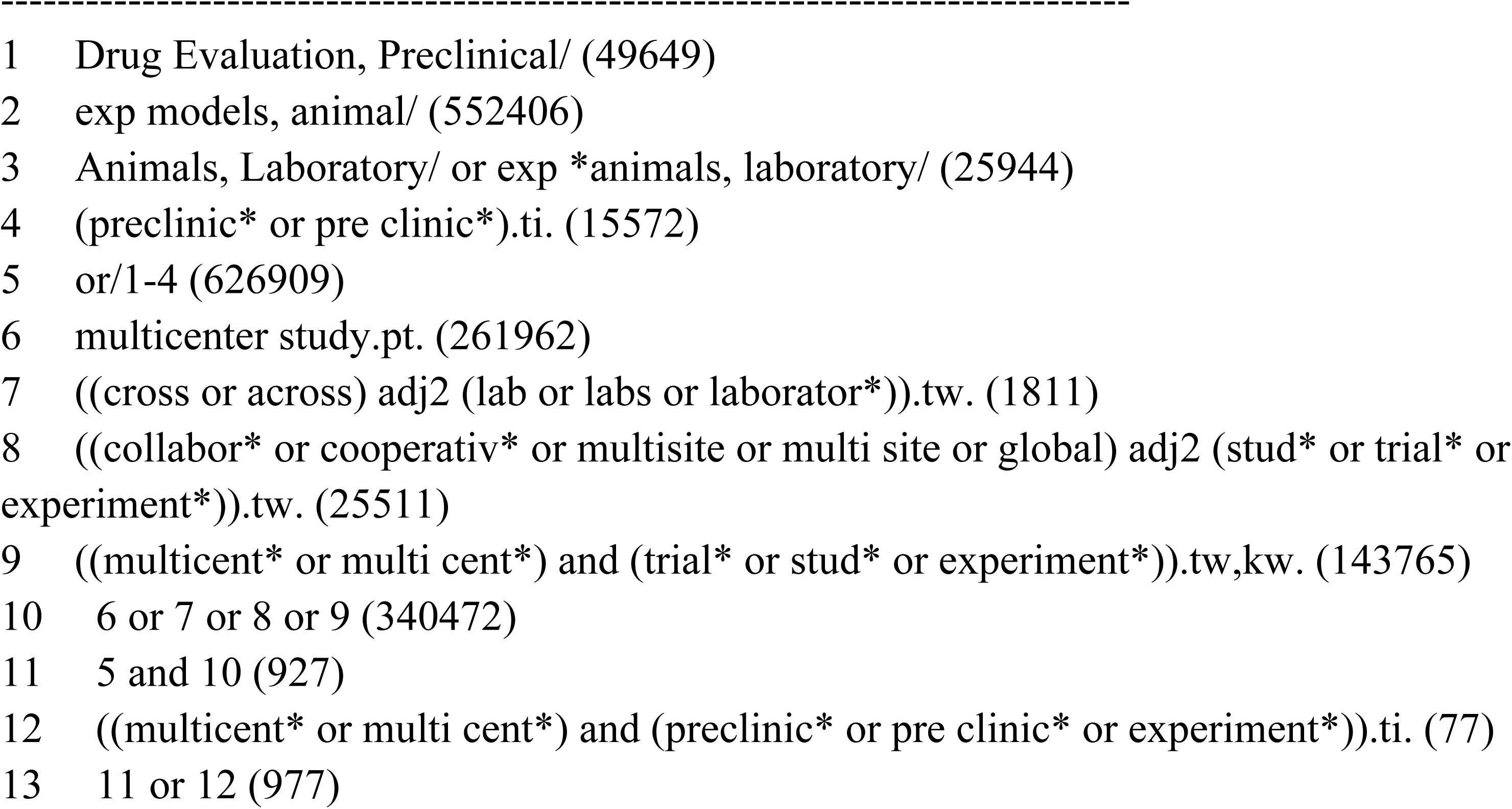

**PEER REVIEW ASSESSMENT: THIS SECTION TO BE FILLED IN BY THE REVIEWER**

Reviewer: Sascha Davis

Date Completed:2018-01-242018-01-24

**Translation**

A- No RevisionA- No Revision

**If “B’ or “C”, please provide and explanation or example**

Click or tap here to enter text.

**Boolean and Proximity Operators**

A- No RevisionA- No Revision

**If “B’ or “C”, please provide and explanation or example**

Click or tap here to enter text.

**Subject Headings**

A- No RevisionA- No Revision

**If “B’ or “C”, please provide and explanation or example**

Click or tap here to enter text.

**Text Word Searching**

A- No RevisionA- No Revision

**If “B’ or “C”, please provide and explanation or example**

Click or tap here to enter text.

**Spelling, Syntax and Line Numbers**

A- No RevisionA- No Revision

**If “B’ or “C”, please provide and explanation or example**

Click or tap here to enter text.

**Limits and Filters**

Choose an item.

**If “B’ or “C”, please provide and explanation or example**

Click or tap here to enter text.

Overall Evaluation

A- No RevisionA- No Revision

Additional Comments: Click or tap here to enter text.

- Could you do “animal model*”.ti or is that too big? Or even “animal adj2 model*”.ti?
- I’m not sure if more wording could be added for the concept of cross-laboratory? – “different labs” or “simultaneous* adj3 laborator*) or (“two laborator* or “three laborator* or “four laborator*”)?

---

**S8 Table.**
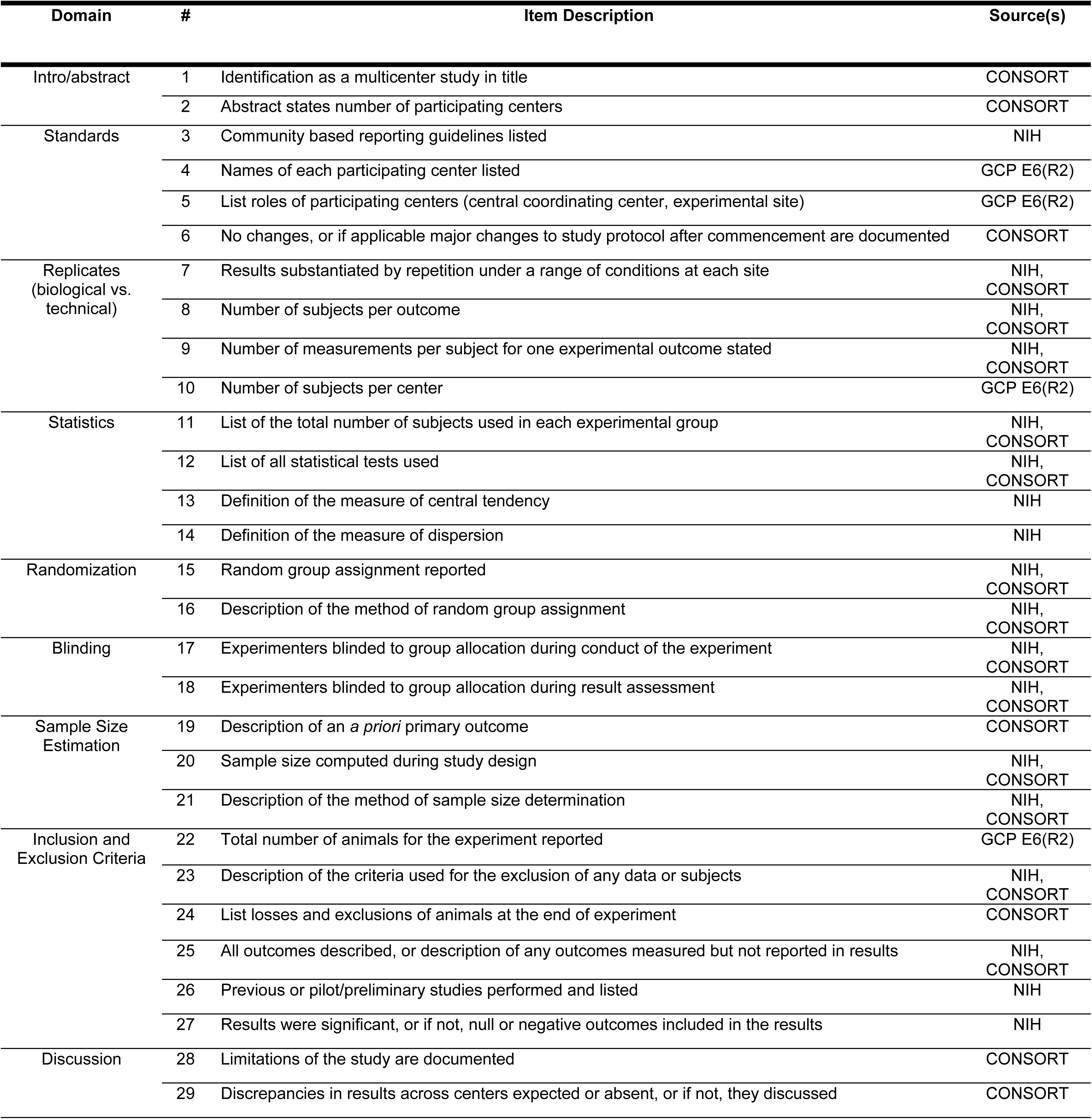
Multicenter reporting checklist with item domain and source(s)

**S9 Table.**
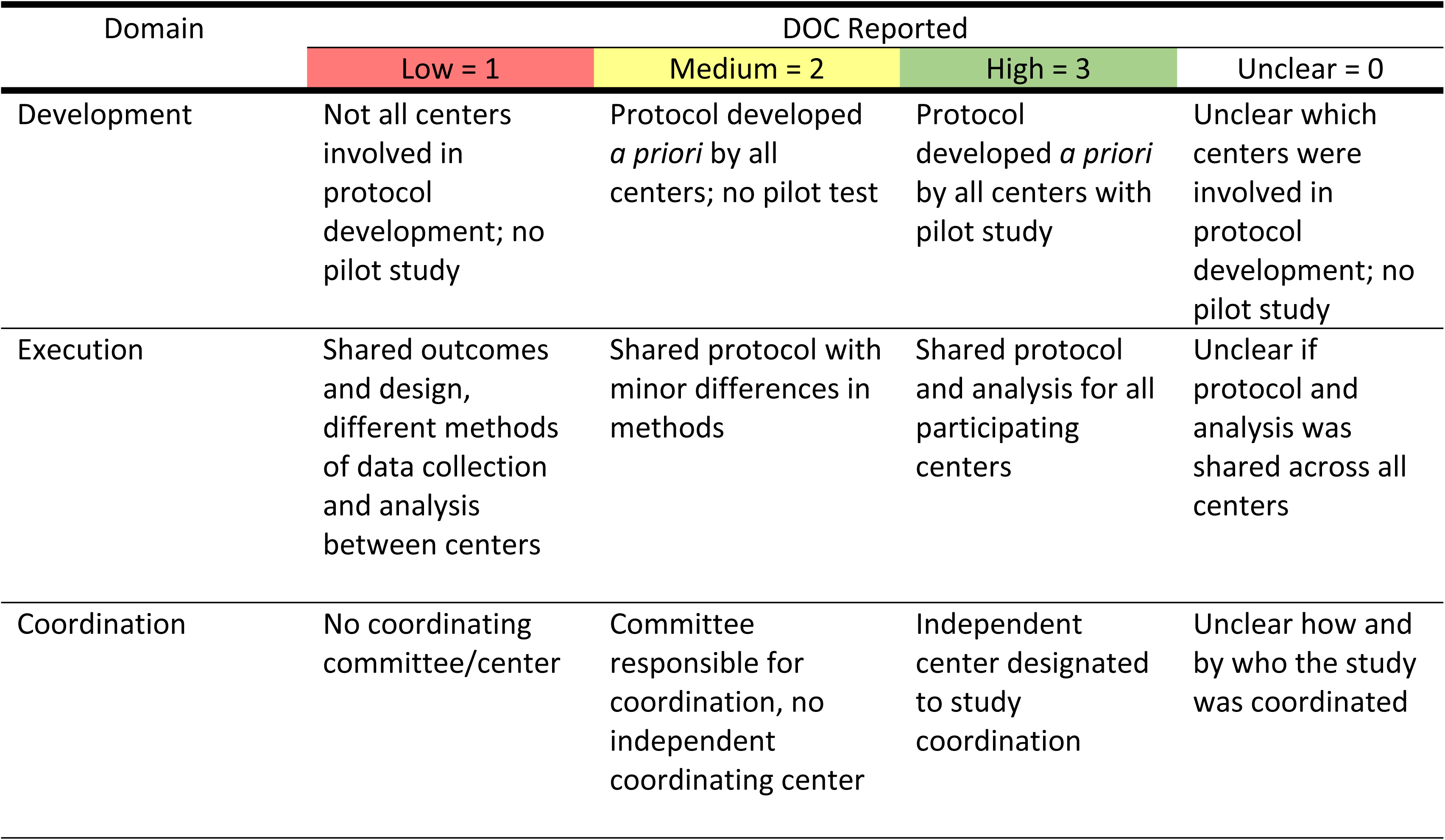
Degree of collaboration assessment criteria for domains

| Domain | DOC Reported |  |  |  |
| --- | --- | --- | --- | --- |
|  | Low = 1 | Medium = 2 | High = 3 | Unclear = 0 |
| Development | Not all centers involved in protocol development; no pilot study | Protocol developed *a priori* by all centers; no pilot test | Protocol developed *a priori* by all centers with pilot study | Unclear which centers were involved in protocol development; no pilot study |
| Execution | Shared outcomes and design, different methods of data collection and analysis between centers | Shared protocol with minor differences in methods | Shared protocol and analysis for all participating centers | Unclear if protocol and analysis was shared across all centers |
| Coordination | No coordinating committee/center | Committee responsible for coordination, no independent coordinating center | Independent center designated to study coordination | Unclear how and by who the study was coordinated |

